# GCN2 mediates access to stored amino acids for somatic maintenance during *Drosophila* ageing

**DOI:** 10.1101/2023.11.14.566972

**Authors:** Joshua N. Johnstone, Christen K. Mirth, Travis K. Johnson, Ralf B. Schittenhelm, Matthew D.W. Piper

## Abstract

Many mechanistic theories of ageing argue that a progressive failure of somatic maintenance, the use of energy and resources to prevent and repair damage to the cell, underpins ageing. To sustain somatic maintenance an organism must acquire dozens of essential nutrients from the diet, including essential amino acids (EAAs), which are physiologically limiting for many animals. In *Drosophila*, adulthood deprivation of each individual EAA yields vastly different lifespan trajectories, and adulthood deprivation of one EAA, phenylalanine (Phe), has no associated lifespan cost; this is despite each EAA being strictly required for growth and reproduction. Moreover, survival under any EAA deprivation depends entirely on the conserved AA sensor GCN2, a component of the integrated stress response (ISR), suggesting that a novel ISR-mediated mechanism sustains lifelong somatic maintenance during EAA deprivation. Here we investigated this mechanism, finding that flies chronically deprived of dietary Phe continue to incorporate Phe into new proteins, and that challenging flies to increase the somatic requirement for Phe shortens lifespan under Phe deprivation. Further, we show that autophagy is required for full lifespan under Phe deprivation, and that activation of the ISR can partially rescue the shortened lifespan of *GCN2*-nulls under Phe deprivation. We therefore propose a mechanism by which GCN2, via the ISR, activates autophagy during EAA deprivation, breaking down a larvally-acquired store of EAAs to support somatic maintenance. These data refine our understanding of the strategies by which flies sustain lifelong somatic maintenance, which determines length of life in response to changes in the nutritional environment.

## Introduction

Ageing is defined as the progressive loss of physiological function over time that leads to death^1^; almost all mechanistic theories of ageing attribute this loss of physiological function to more damage being inflicted upon the cell than is repaired^2–6^. This is thought to be an issue of metabolic resource management, where nutrients are either consumed at insufficient levels to support somatic maintenance, or are in adequate supply, but are prioritised for consumption by other processes (e.g., reproduction) which makes them unavailable to support long life^7^.

There is a middle ground at which point nutritional resources are prioritised for somatic maintenance; reducing levels of food intake to ∼20-50% of the amount eaten by choice is beneficial for lifespan^8^. This moderate dietary restriction (DR) extends lifespan across taxa from yeast to primates^9–13^. Since DR levels of food intake also compromise reproduction^14–16^, it has been assumed that moderate DR extends lifespan by re-prioritising resources away from reproduction and towards life-preserving somatic maintenance^7,17^. While a great deal of research has been devoted to the molecular switches that control resource reallocation^18,19^, little is known about which processes comprise somatic maintenance. One strategy to understand how DR extends lifespan is to define the nutritional requirements for sustained life, and then to characterise the set of somatic processes which are fuelled by these nutrients (i.e., somatic maintenance).

Lifespan extension by DR in *Drosophila melanogaster* and mice can be achieved by reducing the abundance of protein, relative to other macronutrients and micronutrients, in the diet^20–23^. Within an organism, protein causes physiological changes in part via intracellular signalling that is triggered by the abundance of the constituent amino acids (AAs)^24^, and as such considerable effort has been directed to understanding how individual amino acids interact with AA sensors to alter lifespan^25,26^. Recently, it was shown that individually removing each of the essential AAs (EAAs) from the diet of adult *Drosophila* elicits vastly different effects on their lifespan^27^. Specifically, removal of the EAA methionine (Met) reduces lifespan to approximately 25% of fully-fed flies, while removing the EAA threonine (Thr) yields a lifespan approximately 50% that of fully-fed flies. Most strikingly, removal of the EAA phenylalanine (Phe) has no significant impact on lifespan compared to a diet with a full complement of AAs. This variation is surprising because each of these EAAs are strictly required in the diet of *Drosophila* for growth and reproduction^28,29^, suggesting that somatic maintenance processes have different requirements for each individual EAA compared to processes involved in growth and reproduction.

This variation in survival response to each EAA deprivation is entirely dependent on General Control Nonderepressible 2 (GCN2)^27^, an intracellular AA sensor which is conserved from yeast to mammals^30^. GCN2 acts during periods of AA deprivation to repress global translation, slowing growth and preserving AAs, while also activating a specific subset of genes involved in the integrated stress response (ISR)^31–36^. *GCN2*-null flies show a normal lifespan on a complete diet but have equally short lifespans when any one EAA is removed from the diet^27^. Thus, GCN2 serves to protect flies against individual EAA deprivation, but the extent of this protection varies in an AA-specific manner.

Here, we utilise this EAA / GCN2 interaction to interrogate the somatic maintenance processes that sustain adult lifespan in *Drosophila melanogaster*. In particular, we use proteomics, dietary AA manipulations and transgenic interventions to investigate how flies survive without dietary Phe. These data reveal a likely process by which flies store protein during development for later use in somatic maintenance. Future work to characterise this storage and its uses are likely to be important for uncovering the causes of ageing.

## Results

### Phe is unlikely to be provided by commensal bacteria

Recent work has found that Phe can be removed from the diet of adult *Drosophila melanogaster* without imposing any cost to their lifespan^27^. This is surprising because Phe is classified as an essential amino acid and is strictly required for development and reproduction^27,29^.

One explanation for this is that adult flies obtain Phe from a source other than their diet, such as from their gut microbiota. Indeed, dietary depletion of single EAAs can be partially rescued by commensal bacteria supplementation during pre-adulthood – effects that are abolished by treatment with an antibiotic mixture comprising kanamycin, ampicillin, tetracycline and erythromycin^29^. We therefore treated our adult flies with the same antibiotic mixture to investigate whether their ability to survive without Phe is similarly compromised. Antibiotic addition to the diets of adult flies had no effect on lifespan (P = 0.19, ANOVA) whether or not the food contained Phe (P = 0.19) (Figure 1) indicating that commensal or food-borne microbes are unlikely to be supplying Phe to the flies for sustaining lifespan.

**Figure 1.**
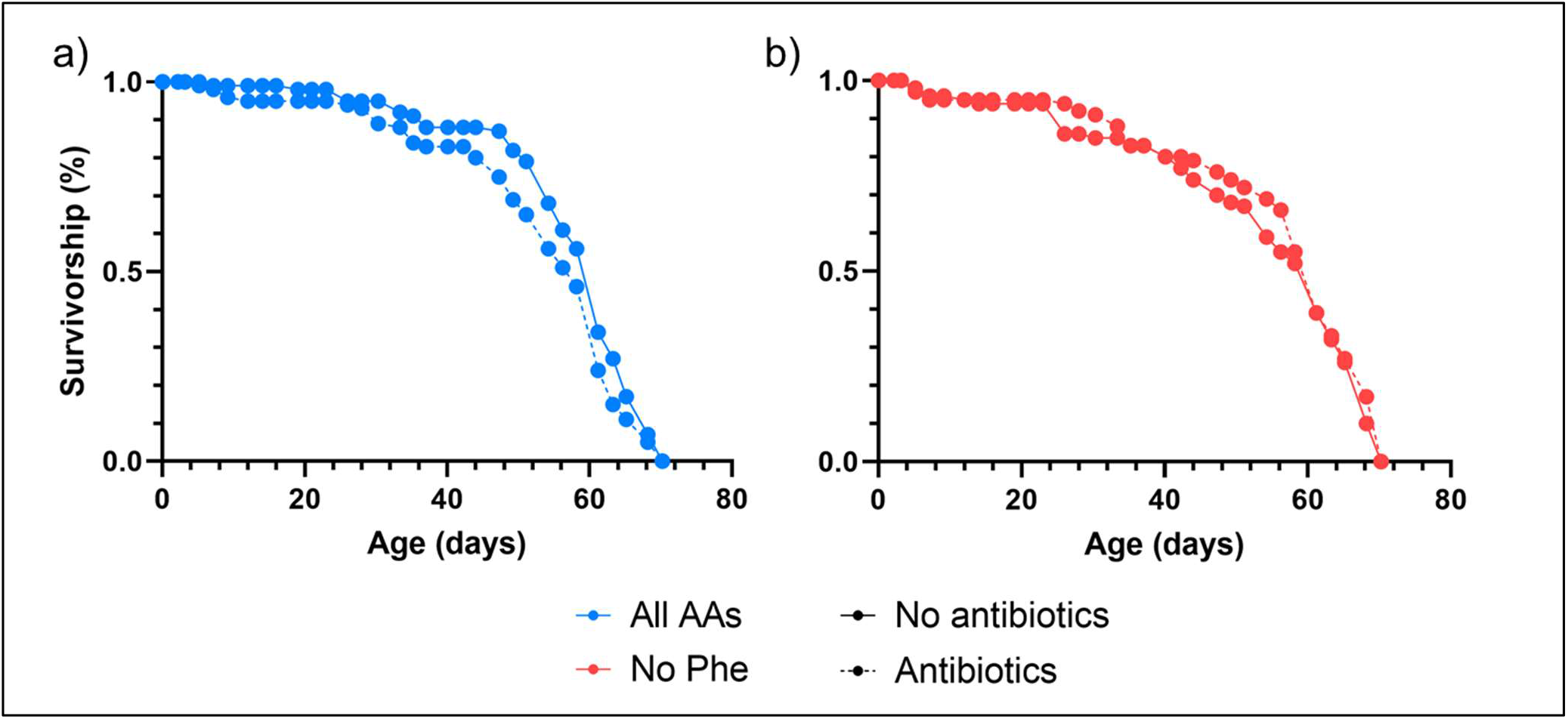
Antibiotic addition has no effect on lifespan under Phe deprivation. To investigate how flies survive when deprived of dietary Phe, we explored whether Phe could be provided by bacteria living in the food media; commensal bacteria have been shown to support *Drosophila* development when EAAs are absent. We added an antibiotic mix to the adult food media which is known to remove these bacteria. a) Wildtype lifespan was not affected by antibiotic addition (P = 0.19, ANOVA) b) nor was there any interaction between antibiotic addition and diet (P = 0.19). These data indicate that it is unlikely that Phe-deprived flies receive Phe from commensal bacteria. Blue lines represent flies fed a full complement of AAs, and red lines represent flies fed a diet missing Phe. Solid lines represent a diet with no antibiotics, and dotted lines represent a diet with antibiotics added.

### Phe-deprived adult *Drosophila* readily incorporate Phe into new proteins

An alternative explanation for why adult flies can survive in the absence of a Phe supply is that they don’t require it; the proteins they express to sustain adult lifespan may require little to no Phe. The fly exome codes for 201 proteins without any Phe, and the remaining proteins contain a Phe content between 0.07% and 20.7%, with an average of 3.7% (FlyBase, 2023).

To investigate the nature of the expressed proteome, we maintained adult *Drosophila* on Phe-free food for 40 days (late middle age) at which point we replaced dietary lysine with a heavy isotope of lysine while maintaining a Phe-free diet. After 4, 9 and 19 days on the lysine-labelled, Phe-free food, we assessed the proteome via mass spectrometry (Figure 2a) to identify newly-synthesised proteins as judged by the incorporation of heavy lysine. We identified a total of 1,749 proteins containing the heavy lysine label at all time points (Table S1). Surprisingly, rather than being low in Phe, these proteins had a significantly higher Phe content than randomly selected, similarly sized groups of proteins encoded by the fly exome (P < 0.001, T-test) (Figure 2b). Thus, despite being deprived of Phe for more than half of their adult lifespan, the flies continue to express a Phe-enriched proteome, suggesting that flies possess a reservoir of Phe that remains accessible for later life protein synthesis.

**Figure 2.**
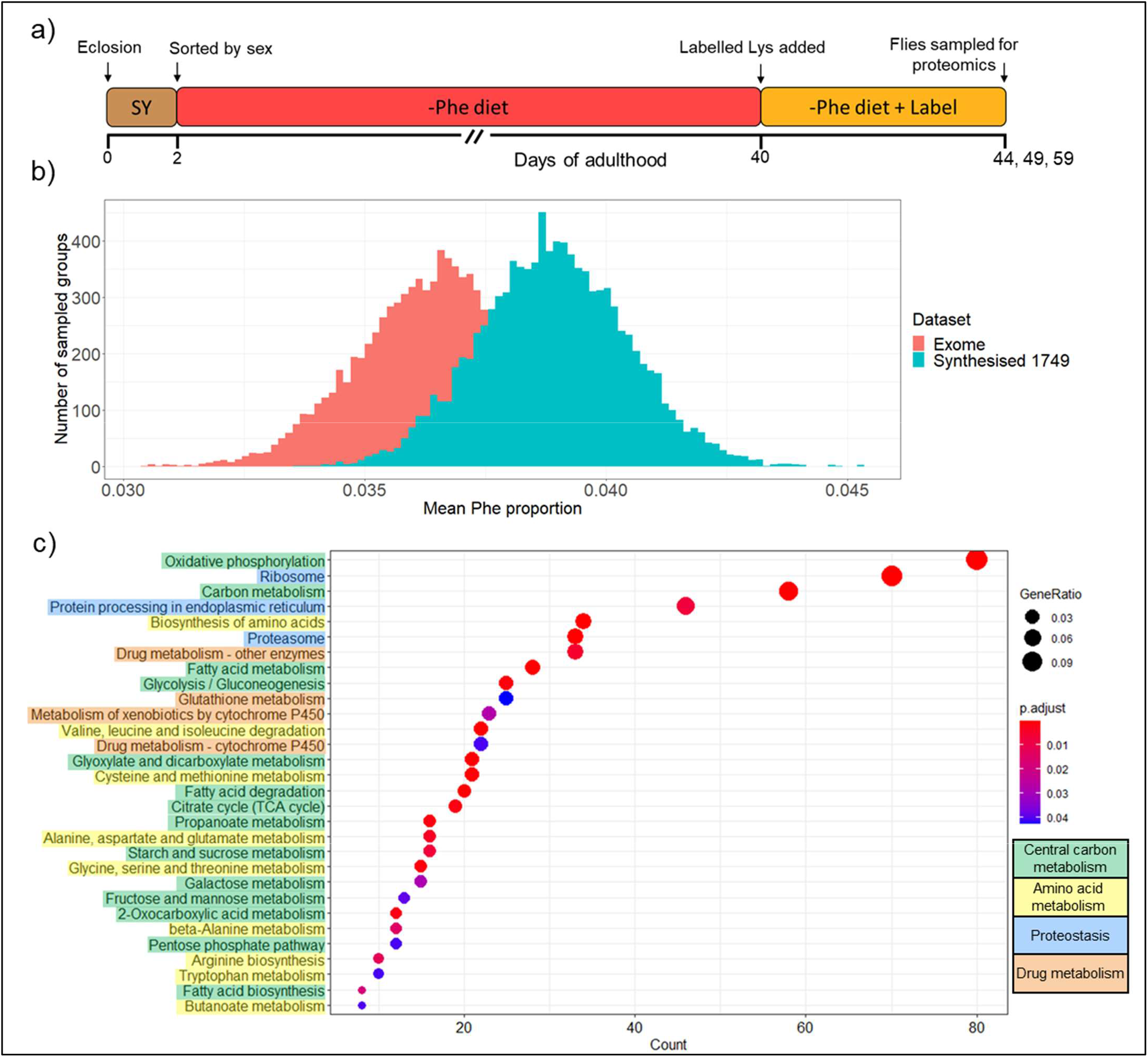
Phe-starved flies continue to synthesise Phe-rich proteins. To determine whether Phe-starved flies continue to use Phe for protein synthesis, a) we starved female adult *Drosophila* of Phe for 40 days before labelling and detecting newly-synthesised proteins by mass spectrometry. b) The 1749 proteins found to be newly-synthesised after 40 days of Phe deprivation were richer in Phe than randomly sampled proteins from the exome (P < 0.001, T-test). c) We performed functional enrichment analysis on the list of 1749 proteins. The size of each dot represents the proportion of the total protein set which belongs to each respective gene ontology category. The colour of the dot represents the probability of achieving the observed proportions by randomly sampling the proteome for the same sized sets of proteins. Proteins related to central carbon metabolism, amino acid metabolism and translation were found to be over-represented, as were detoxification proteins, which made up >14% of all proteins identified in Phe-deprived flies.

As this set of proteins continues to be expressed even after extended EAA deprivation, we reasoned that this set of proteins may be particularly important for sustaining life. Functional enrichment analysis of the heavy lysine-containing proteins revealed functional over-representation of central carbon metabolism (in particular metabolism of fatty acids, as well as sugars through glycolysis, the pentose phosphate pathway, the TCA cycle and oxidative phosphorylation), AA metabolism, and proteostasis (including various AA metabolic pathways, protein breakdown via the proteasome, protein synthesis due to ribosome expression, and protein processing via the ER) (p < 0.05) (Figure 2c). In addition to these core cellular functions, proteins related to the removal of xenobiotic (foreign) and endogenous metabolic toxins were also over-represented, comprising >14% of all proteins detected. It is therefore possible that during Phe deprivation, flies selectively sustain the expression of proteins with functions related to core metabolic pathways, as well as functions related to detoxification capacity, and these represent the core set of proteins required to sustain life and protect against molecular damage accumulation.

### Phe deprivation does not protect against Thr deprivation

If chronic Phe deprivation acts as a specific trigger to activate lifespan-preserving processes, perhaps Phe deprivation can protect fly lifespan against simultaneous deprivation of another EAA that would normally shorten lifespan. To test this, we combined dietary Phe deprivation with dietary Thr deprivation, where Thr deprivation alone shortens lifespan to about 50% of fully fed controls^27^. However, rather than ameliorating the cost to lifespan imposed by Thr deprivation, concurrent deprivation of Phe with Thr slightly worsened this cost and reduced lifespan (P = 0.002, Tukey’s HSD) (Figure 3a).

**Figure 3.**
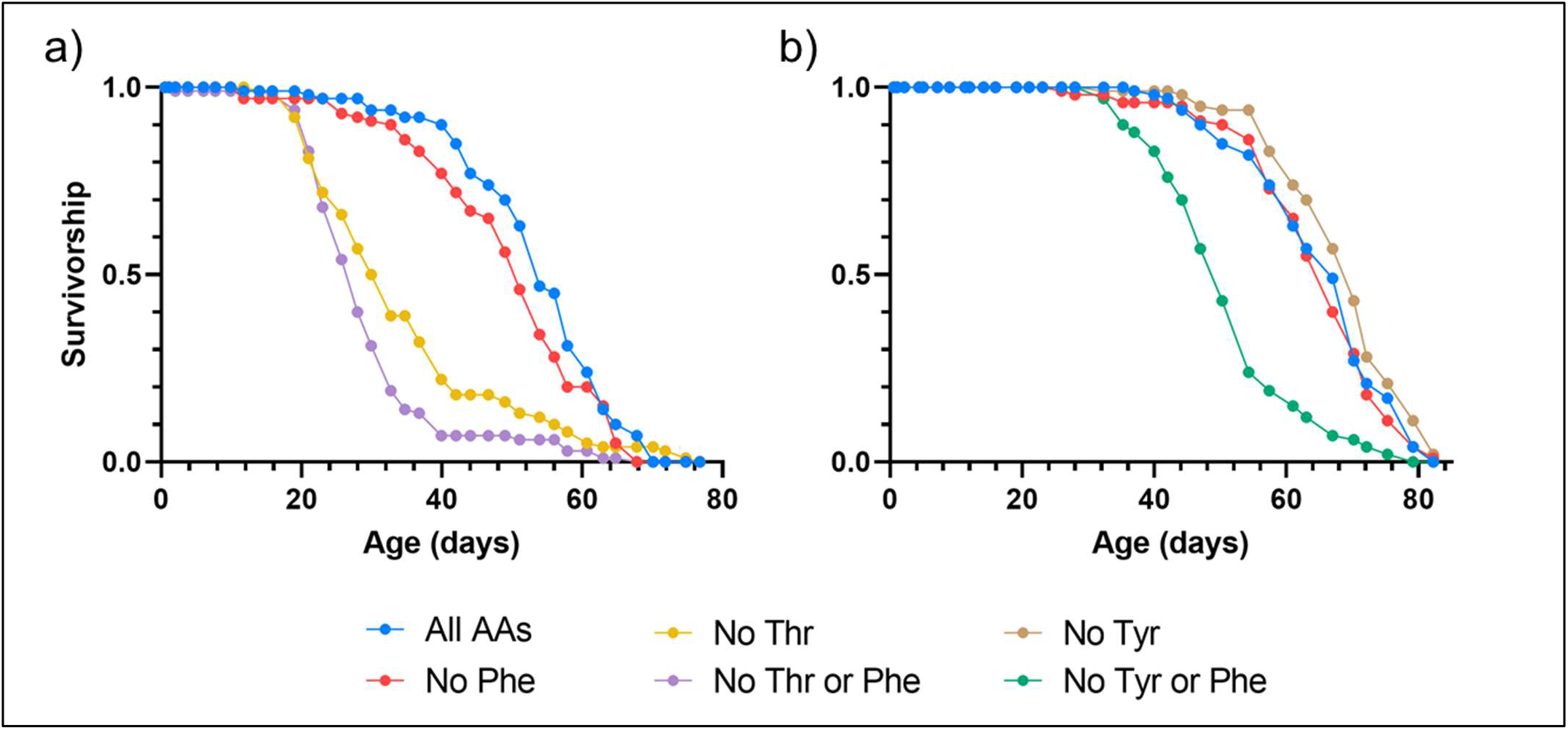
Lifespan is limited by insufficient supply of individual EAAs relative to demand. Given that Phe deprivation is the only EAA deprivation with no lifespan cost, we asked whether Phe deprivation could protect against other EAA deprivations. a) We deprived adult female *Drosophila* of both Phe and another EAA Thr. Wildtype flies deprived of both Phe and Thr were shorter lived than those deprived of only Thr (P = 0.002, Tukey’s HSD), indicating that Phe deprivation cannot protect against Thr limitation. To test whether increasing the physiological demand for Phe can limit lifespan, b) we deprived adult female *Drosophila* of both Phe and Tyr, which is synthesised from Phe. Wildtype flies simultaneously deprived of Phe and Tyr were indeed shorter lived than flies fed all AAs (P < 0.001) or deprived of Phe (P < 0.001) or Tyr (P < 0.001) alone. These data suggest that lifespan is limited by insufficient supply of individual EAAs relative to their somatic maintenance demand.

One explanation for this failure of Phe deprivation to rescue Thr deprivation is that sustained life under dietary EAA deprivation requires access to internally stored EAAs, and lifespan will be limited if these EAAs are not stored or cannot be accessed in sufficient amounts. In this case, the flies’ pool of stored Thr is relatively small compared to the amount required to sustain life, whereas the pool of stored Phe is sufficiently large to sustain protein synthesis for a full lifespan (as supported by our proteomics data). Thus, combinations of EAA restrictions may limit lifespan to a level that can only be sustained to the extent supported by the most limiting EAA.

### Increasing the requirement for Phe reduces lifespan under Phe deprivation

If each individual EAA deprivation limits lifespan according to an intrinsic relationship between its level accessible from stores and the specific requirement for that EAA to support life, it should be possible to shorten the lifespan of Phe-deprived flies by accelerating the depletion of their Phe reservoir. In *Drosophila,* the non-essential AA (NEAA) tyrosine (Tyr) is synthesised from Phe via an irreversible reaction catalysed by phenylalanine hydroxylase^37^. Thus, maintaining flies on food lacking both Phe and Tyr may shorten lifespan because Phe will be depleted more rapidly to meet the need for both Phe and its catabolite Tyr. By contrast, depriving flies of either Phe or Tyr alone should not impose a cost to lifespan because sufficient stores of Phe exist to meet the need for each one individually.

Consistent with our storage hypothesis, removing Phe or Tyr alone resulted in lifespans that were no different from fully fed controls (P > 0.170, Tukey’s HSD), whereas depriving flies of both Phe and Tyr resulted in a 25% reduction in lifespan when compared to flies fed all AAs (P < 0.001) (Figure 3b). Together, these data indicate that while Phe is usually stored in sufficient amounts to sustain lifelong somatic maintenance, this storage can become limiting when somatic maintenance requirements for Phe are increased. This further supports our hypothesis that lifespan under each EAA deprivation is determined by a balance between the amount of the EAA stored and the amount required for somatic maintenance.

### Alternative activation of nutritional stress signalling partially restores the lifespan of *GCN2*-null flies under Phe deprivation

Recent data has shown that the ability of flies to maintain a full lifespan without dietary Phe requires the presence of the AA sensor and signalling kinase GCN2^27^. When *GCN2*-null flies are fully fed, they are equally as long lived as fully-fed controls, but when deprived of dietary Phe, their lifespan drops to about 30% of wild type flies maintained with or without Phe (P < 0.001 for all comparisons of Phe-deprived *GCN2*-nulls versus controls, Tukey’s HSD) (Figure 4a). When activated by AA deprivation, GCN2 initiates the integrated stress response (ISR), which involves repression of general translation as well as coordinating enhanced expression of a subset of genes involved in protecting the flies against nutrient stress^31,33–36^. *GCN2*-null flies may therefore be protected against Phe deprivation by alternative activation of the ISR.

**Figure 4.**
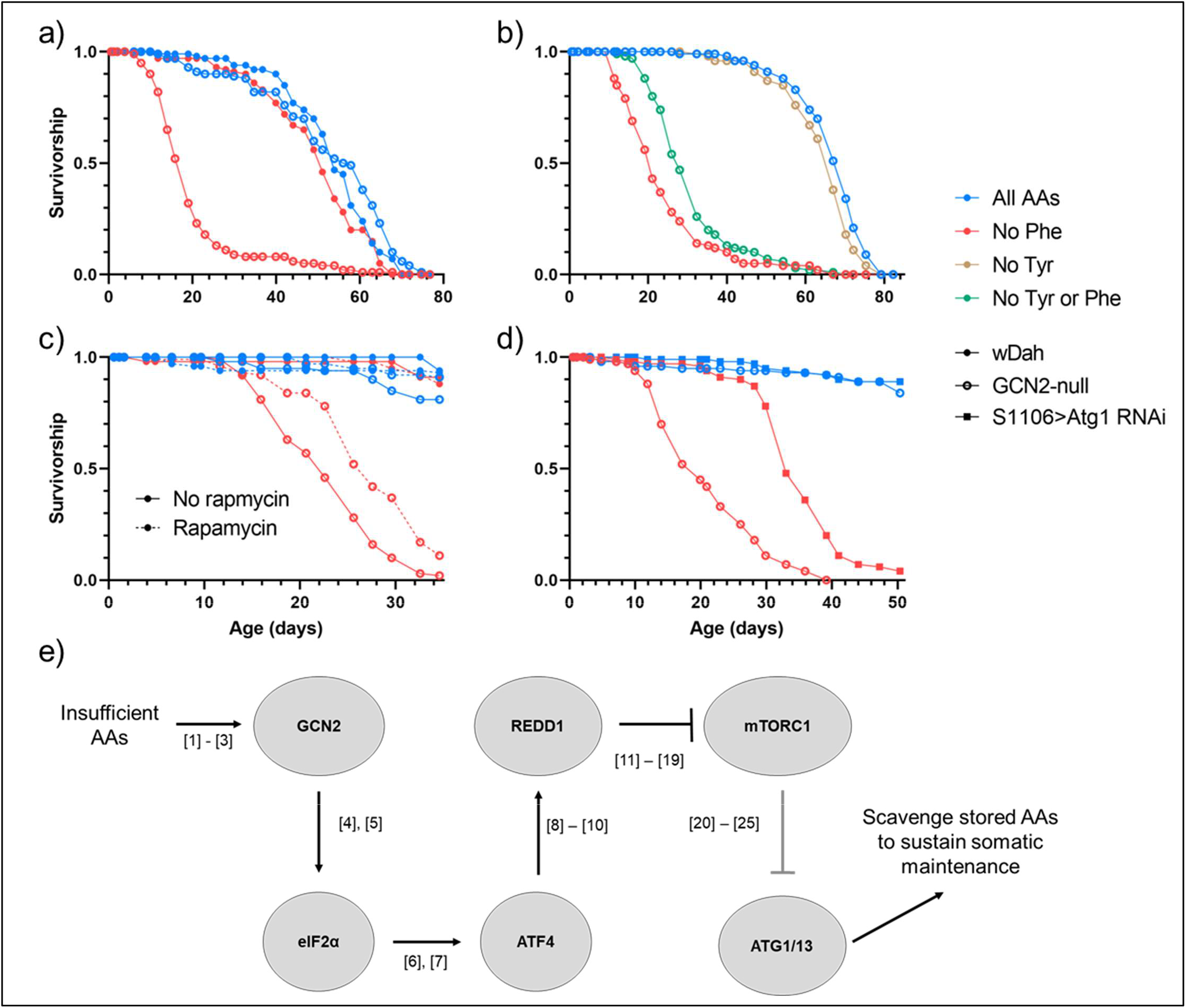
GCN2-mediated autophagy may release stored EAAs under EAA deprivation. a) Starving *GCN2*-null flies of Phe shortens lifespan in comparison to *GCN2*-null flies fed all AAs and wildtype flies fed all AAs or no Phe (P < 0.001 in all comparisons). b) To test whether lifespan of GCN2-null flies under Phe deprivation can be rescued by ISR activation, we deprived flies of both Phe and Tyr and measured lifespan. Deprivation of Tyr partially rescued GCN2-null lifespan from the cost of Phe deprivation (P = 0.017). c) The ISR can also be activated via pharmacological inhibition of mTOR. Addition of the mTOR inhibitor rapamycin to the food partially rescues GCN2-null lifespan under Phe deprivation (P < 0.001). d) ISR activation ultimately results in an upregulation of autophagy. To test whether autophagy is required for survival under Phe deprivation, we expressed Atg1 RNAi in the fat body and gut then deprived the flies of Phe. Atg1 RNAi flies deprived of Phe had reduced lifespan compared to those fed all AAs (P < 0.001). e) These lifespan data suggest that under EAA deprivation, GCN2 may activate autophagy to release stored EAAs and thus sustain lifelong somatic maintenance. Our proposed model for this mechanism combines many individually well-characterised signalling events, from studies in an array of model systems, to form a pathway that connects AA deprivation to autophagy and encompasses the two major AA sensors (GCN2 and mTOR). Grey ovals represent proteins involved in the signalling pathway. Each protein interaction is accompanied by a reference in square brackets for a study in which the interaction has been observed, and these interactions are expanded upon in Table S2.

Recent work has shown that Tyr deprivation can activate the ISR independently of GCN2^38^, meaning that Tyr deprivation should protect *GCN2*-null flies against Phe deprivation by restoring activation of the ISR. In line with this prediction, *GCN2*-null flies deprived of both Phe and Tyr lived 33% longer than *GCN2*-nulls deprived of Phe alone (P = 0.017) (Figure 4b). While this is a significant enhancement of lifespan, it is far from complete restoration. This may occur if Tyr deprivation only partially activates the ISR, or because Tyr deprivation can only provide a limited benefit for lifespan; our previous data with wild type flies show that its absence becomes deleterious later in life when Phe is also missing from the diet (Figure 3b).

The mild protective effect of Tyr depletion for *GCN2*-nulls may be caused by activation of the ISR to break down AA stores for the synthesis of new proteins, and/or by sparing the flies from investing in the proteome by generally suppressing translation. Both of these effects can also be achieved by inhibiting the AA sensor and growth activator mTORC1^39–44^. We therefore added the mTOR inhibitor rapamycin to the food of the flies to see if mTORC1 suppression would rescue the lifespan of *GCN2-*nulls on food without Phe. Rapamycin addition extended the lifespan of *GCN2*-nulls on Phe-free food by 21% (P < 0.001, Tukey’s HSD) and, to the extent that we measured it, had no effect on the lifespan of wildtype flies when fed diets with all AAs or lacking Phe, nor did it affect the lifespan of *GCN2*-nulls fed all AAs (P > 0.12 in all comparisons) (Figure 4c).

### Autophagy knockdown reduces lifespan under Phe deprivation

If retrieval of AAs from protein stores is a key element of these lifespan effects, then activation of autophagy may be required for flies to survive EAA deprivation. To test this, we knocked down the autophagy protein ATG1 specifically in the fly fat body and gut^45,46^, which are major metabolic tissues and are known to be important reservoirs for nutrients^47,48^. *Atg1* knockdown significantly shortened lifespan of flies on food without Phe (P < 0.001), but not to the same extent as caused by loss of *GCN2* (P < 0.001) (Figure 4d). By contrast, *Atg1* RNAi flies were substantially longer lived when maintained on a complete diet and, to the extent of our experiment, were no different from fully fed *GCN2*-null flies (P = 0.39).

Together, our data point to flies possessing a developmentally acquired store of EAAs, which they access during dietary EAA deprivation by activation of the ISR to trigger autophagy. The EAAs liberated by this process are then utilised for the synthesis of proteins required for lifelong somatic maintenance.

## Discussion

To sustain life, an organism must employ a specific set of processes required for physiological function and ongoing somatic maintenance. Sustaining these processes requires resources that are originally sourced from the diet. Because these resources are generally thought to be limiting, the way in which they are differentially allocated between fitness traits lies at the heart of explaining the constraints under which ageing has evolved^17,49–51^. In seeking mechanistic explanations for ageing, many studies have focused on defining the nutritional elements that limit organismal fitness; nitrogen limitation, which for flies takes the form of limitation for an essential AA, has recently become a central focus of these studies^20–22^. Here, our data, alongside those of Srivastava *et al.*^27^, indicate how limiting amounts of dietary AAs determine adult *Drosophila* lifespan. Overall, our data are consistent with a model in which flies store AAs during development, which they can later access to protect themselves while they find a better food source to support reproduction. We propose that the ability to sustain life under each EAA deprivation could be a read out of the extent to which an EAA is stored relative to the degree to which it is required for somatic maintenance.

These data shed light on the mechanism used by flies to protect themselves during the period of time between experiencing an EAA/total protein deficit and finding a proteinaceous food source. Deprivation of any AA is thought to result in a build-up of uncharged tRNA molecules, which directly bind to and activate GCN2^31,32,52,53^. Given that *GCN2*-null flies quickly succumb to any EAA deprivation, GCN2 may be responsible for providing access to stored EAAs for sustained somatic maintenance during EAA deprivation. Our data indicate that this breakdown occurs via autophagy, the process of degrading and recycling cellular components, which is the most prominent route for bulk AA recycling^54^.

Evidence from a variety of sources and study systems indicate that GCN2 activation can lead to autophagy induction via mTORC1 inhibition, and therefore the breakdown of stored protein (Figure 4e and Table S2). Under this mechanism, GCN2 activation enhances the translation of selected transcripts including that of the transcription factor ATF4^31,33–36^. While there is currently no evidence that GCN2 or ATF4 can activate autophagy directly, evidence from murine cell culture indicates that ATF4 upregulates *REDD1*, which encodes a kinase known to suppress the AA sensor complex mTORC1^55,56^, in turn activating autophagy^39–44^. This is consistent with our data in which Phe deprivation activates GCN2, which indirectly induces autophagy to liberate stored EAAs for ongoing survival and somatic maintenance.

Given the proposed role of GCN2 in freeing stored EAAs to support lifespan under EAA deprivation, the variability in lifespan response to each EAA deprivation can be explained by the difference between the amount of that EAA that is stored, and the amount that is required for somatic maintenance. In our study, we focused on Phe deprivation since, unlike adulthood deprivation of any one of the other EAAs, omitting Phe from the adult diet has no effect on wildtype fly lifespan^27^. This indicates that Phe is atypical among the EAAs in that it is stored in excess of its requirement for lifelong somatic maintenance. By contrast, dietary Thr deprivation shortens lifespan considerably and dietary Met deprivation shortens it to an even greater extent^27^. By our model, the stored amounts of these EAAs fall far short of the levels required for flies to sustain lifelong somatic maintenance.

One of the proteins found in our proteomics data to be continuously synthesised in flies despite suffering from chronic Phe-deprivation is larval serum protein 2 (LSP2), which is strongly induced during late larval development and is particularly rich in Phe^57^. It therefore represents a candidate protein to store larvally acquired Phe that can be used to fuel ongoing somatic maintenance. Knockdown of *Lsp2* in developing flies should reduce this store and therefore reduce lifespan under adult Phe deprivation.

Our proteomics data can also reveal the nature of the processes that underpin somatic maintenance. We found an over-representation of proteins involved in central carbon metabolism, proteostasis and cellular detoxification. These proteins may represent the baseline requirement for keeping flies alive. If it is indeed true that fuelling cellular detoxification is required to keep cellular damage in check to sustain life^6^, knockdown of cap-n-collar (*cnc*) (the *Drosophila* homolog of the xenobiotic response transcription factor Nrf2) should reduce activity of the cellular detoxification processes and shorten lifespan under Phe deprivation. Such a response has been observed in *C. elegans*, where loss of the *Nrf2* ortholog *skn-1* shortens lifespan, and *skn-1* overexpression increases lifespan^58,59^.

## Conclusions

In this study, by investigating the relationship between EAA deprivation, GCN2, and lifespan, we identify EAA storage and retrieval as a likely key regulator of adult *Drosophila* lifespan. We point to a new approach for identifying the resources, and molecular processes, that are required to sustain life. Future studies that identify and characterise these processes are likely to enhance our understanding of the mechanisms that modulate ageing.

## Methods

### Fly husbandry

An outbred, white-eyed Dahomey strain of *Drosophila melanogaster* (wDah) was used for all experiments unless otherwise specified. The *GCN2*-null flies used were obtained as a gift from Dr. Sebastian Grönke (Max Planck Institute for Biology of Ageing). The S1106-GeneSwitch driver line^45^ was obtained as a gift from Prof. Linda Partridge (University College London). The UAS-Atg1-RNAi line^46^ was obtained as a gift from Dr. Donna Denton (University of South Australia). All flies were reared at a controlled population density using eggs laid by age-matched mothers, as described by Linford *et al*.^60^, and developed to adulthood on a standard sugar-yeast (SY) medium as described by Bass *et al*.^61^. Newly emerged adult flies were allowed to mate for two days on the SY medium to standardise the mating status of all flies, then 48 h old adult female flies were sorted under light CO_2_ anaesthesia (< 30 minutes) and placed onto their respective experimental diet. During both rearing and experimental stages flies were kept in a controlled environment of 25°C temperature, 70% humidity and a 12-hour light/dark cycle. wDah stocks were maintained at 25°C temperature, 70% humidity and a 12-hour light/dark cycle while *GCN2-*null, S1106-GeneSwitch and UAS-Atg1-RNAi stocks were maintained at 18°C, 60% humidity and a 12-hour light/dark cycle.

### Diets

Flies were developed to adulthood on a standard sugar-yeast (SY) medium as described by Bass *et al*.^61^. Chemically-defined synthetic (holidic) diets were prepared as described by Piper *et al*.^62,63^, following the exome-matched FLYAA formula. Where antibiotics were used, an antibiotic mix was prepared as described by Consuegra *et al*.^29^, consisting of kanamycin (50 μg/mL), ampicillin (50 μg/mL), tetracycline (10 μg/mL) and erythromycin (5 μg/mL). Stock solutions of each antibiotic were made in either MilliQ water (ampicillin and kanamycin) or ethanol (erythromycin and tetracycline) based on their solubility, such that adding of 1mL stock to 1L holidic media resulted in the above final concentrations. Immediately before dispensing the media, each antibiotic solution was added with a micropipette, under constant stirring to promote even mixing. A separate length of tubing was used to dispense the food containing antibiotics, ensuring no antibiotics were inadvertently introduced to the antibiotic free media. Where rapamycin was used, 30 μL absolute EtOH (Sigma-Aldrich) containing 1000 μM rapamycin (Sigma-Aldrich) was pipetted on top of 3 mL of holidic food for a final rapamycin concentration of 10 μM. We added 30 μL of EtOH to control vials. The food was then left at room temperature for 24 hours to allow the drug to disperse into the food before use. Where the S1106-GeneSwitch driver was used, 30uL absolute EtOH (Sigma-Aldrich) containing 1.67 mM RU486 (Sigma-Aldrich) was pipetted on top of 2 mL of holidic food for a final RU486 concentration of 25 μM. Control vials received 30 μL of EtOH. The food was then left at room temperature for 24 hours to allow the drug to disperse into the food before use.

### Proteomics

Adult female flies were starved of Phe for 40 days, then the lysine in the food was completely replaced by a heavy isotope of lysine in which all 6 carbons are 13C (Silantes). After 4, 9 and 19 days of exposure to the label, 10 flies per condition were sampled and snap-frozen in liquid nitrogen. Protein was extracted from the samples using sodium deoxycholate (SDC) solubilization essentially as described previously^64^. Using a Dionex UltiMate 3000 RSLCnano system equipped with a Dionex UltiMate 3000 RS autosampler, an Acclaim PepMap RSLC analytical column (75 µm x 50 cm, nanoViper, C18, 2 µm, 100Å; Thermo Scientific) and an Acclaim PepMap 100 trap column (100 µm x 2 cm, nanoViper, C18, 5 µm, 100Å; Thermo Scientific), the tryptic peptides were separated by increasing concentrations of 80% acetonitrile (ACN) / 0.1% formic acid at a flow of 250 nl/min for 158 min and analyzed with an Orbitrap Fusion mass spectrometer (ThermoFisher Scientific). The instrument was operated in data dependent acquisition mode to automatically switch between full scan MS and MS/MS acquisition. Each survey full scan (m/z 375–1575) was acquired with a resolution of 120,000 (at m/z 200) in the Orbitrap after accumulating ions with a normalized AGC (automatic gain control) target of 1e6 and a maximum injection time of 54 ms. The 12 most intense multiply charged ions (z ≥ 2) were sequentially isolated and fragmented in the collision cell by higher-energy collisional dissociation (HCD) with a resolution of 30,000, an AGC target of 2e5 and a maximum injection time of 54 ms. Dynamic exclusion was set to 15 seconds.

The raw data files were analysed with the MaxQuant software suite v1.6.2.10^65^ and its implemented Andromeda search engine^66^ to obtain protein identifications using a Uniprot Drosophila database downloaded in September 2018. Carbamidomethylation of cysteine residues was selected as fixed modification, whilst oxidation of methionine and acetylation of protein N-termini were set as variable modifications. Up to 3 missed cleavages were permitted, the multiplicity was set to 2 considering Lys6 as heavy labels and a false discovery rate (FDR) of 1% was allowed for both protein and peptide identification. The proteomics data were further analysed with Perseus v1.6.2.3^67^.

### Functional profile analysis

The list of these labelled (newly synthesised) proteins was imported to R Studio^68^, and the “enrichKEGG” and “dotplot” functions of the “clusterProfiler” package^69^ were used for functional profile analysis and visualisation, respectively. To create the distribution of mean Phe content of the proteins, 10000 groups of 100 random proteins were sampled from both the list of 1749 heavy lysine-labelled proteins and from the fly exome (obtained from FlyBase^70^), and the mean Phe content of each group of 100 proteins was plotted.

### Lifespan experiments

After being raised and allowed to mate on SY food, 48 h old adult female flies were sorted under light CO_2_ anaesthesia (< 30 minutes) and placed onto the various diets at a density of 10 females per vial. From this day the protocol of Linford *et al*.^60^ was followed, with flies transferred to fresh food every 2 days and any deaths or censors recorded using the dLife software^60^. Constantly providing fresh food decreased the likelihood of larvae hatching and digging in the food, which can cause the adult flies to drown. Flies that drowned or escaped during food changes were censored at that time point.

## Statistical analyses

### Functional profile analysis

Over-representation of proteins from specific gene ontology categories was determined using the ‘enrichKEGG’ function of the ‘clusterProfiler’ package^69^, using a strict adjusted p-value cutoff of 0.05. Significant values identified groups of proteins that were over-represented in the protein set, as compared to their representation in the *Drosophila* genome sourced from the Kyoto Encyclopedia of Genes and Genomes^71^ (KEGG, organism code = “dme”, accessed 09/02/2021).

### Lifespan data

Raw data generated by the dLife software^60^ was analysed in R Studio^68^, using the Cox Proportional-Hazards Model (‘coxph’) function of the “Survival” package^72^ to find significant differences between genotypes and diets. The ANOVA (‘Anova’) function of the “car” package^73^ was used to assess genotype by diet interactions, and the ‘contrast’ and ‘emmeans’ functions from the “emmeans” package^74^ were used for post-hoc pairwise comparisons (Lenth, 2021).

## Author contributions

J.N.J., M.D.W.P., C.K.M., and T.K.J. conceived the project. R.B.S. performed the mass spectrometry runs and analysed the raw data. J.N.J. performed the other experiments and analysed the data. J.N.J. wrote the original manuscript, and M.D.W.P., C.K.M., and T.K.J. assisted in reviewing and editing. M.D.W.P., C.K.M., and T.K.J. supervised the project. All authors approved the final manuscript.

## Acknowledgements

We thank Amy Dedman and Mia Wansbrough for their help with data collection, especially during the COVID-19 lockdown restrictions of 2020 and 2021. We also acknowledge Tahlia Fulton for their help with backcrossing lines and maintaining fly stocks. This work was funded by a National Health and Medical Research Council grant awarded to M.D.W.P. and T.K.J. (no. APP1182330), Australian Research Council grants awarded to M.D.W.P. (no. FT150100237) and C.K.M. (FT170100259), and a National Institute of Health grant awarded to T.K.J. (no. 5U01HG007530-08). J.N.J. is supported by an Australian Government Research Training Program (RTP) scholarship. This study used BPA-enabled (Bioplatforms Australia) / NCRIS-enabled (National Collaborative Research Infrastructure Strategy) infrastructure located at the Monash Proteomics and Metabolomics Platform.

## Supplementary tables

**Table S1 –** List of proteins synthesised by Phe-deprived flies

**Table S2 –** Protein interactions as referenced in Figure 4

## References

1 López-Otín C, Blasco MA, Partridge L, Serrano M, Kroemer G. The Hallmarks of Aging. Cell 2013;153:1194–217. 10.1016/j.cell.2013.05.039.

2 Weismann A. Ueber die Dauer des Lebens. Am J Psychology 1882;3:105. 10.2307/1411530.

3 Harman D. Aging: A Theory Based on Free Radical and Radiation Chemistry. J Gerontology 1956;11:298–300. 10.1093/geronj/11.3.298.

4 Gensler HL, Bernstein H. DNA Damage as the Primary Cause of Aging. Q Rev Biology 1981;56:279–303. 10.1086/412317.

5 Bjorksten J. The crosslinkage theory of ageing. J Am Geriatr Soc 1968;16:408–27. 10.1111/j.1532-5415.1968.tb02821.x.

6 Gems D, McElwee JJ. Broad spectrum detoxification: the major longevity assurance process regulated by insulin/IGF-1 signaling? Mech Ageing Dev 2005;126:381–7. 10.1016/j.mad.2004.09.001.

7 Kirkwood TBL, Holliday R. The evolution of ageing and longevity. Proc R Soc Lond Ser B Biol Sci 1979;205:531–46. 10.1098/rspb.1979.0083.

8 Weindruch R, Walford RL, Fligiel S, Guthrie D. The Retardation of Aging in Mice by Dietary Restriction: Longevity, Cancer, Immunity and Lifetime Energy Intake. J Nutrition 1986;116:641–54. 10.1093/jn/116.4.641.

9 McCay CM, Crowell MF, Maynard LA. The Effect of Retarded Growth Upon the Length of Life Span and Upon the Ultimate Body Size. J Nutrition 1935;10:63–79. 10.1093/jn/10.1.63.

10 Klass MR. Aging in the nematode Caenorhabditis elegans: Major biological and environmental factors influencing life span. Mech Ageing Dev 1977;6:413–29. 10.1016/0047-6374(77)90043-4.

11 Chapman T, Partridge L. Female fitness in *Drosophila* melanogaster: an interaction between the effect of nutrition and of encounter rate with males. Proc Royal Soc Lond Ser B Biological Sci 1996;263:755–9. 10.1098/rspb.1996.0113.

12 Jiang JC, Jaruga E, Repnevskaya MV, Jazwinski SM. An intervention resembling caloric restriction prolongs life span and retards aging in yeast. Faseb J 2000;14:2135–7. 10.1096/fj.00-0242fje.

13 Colman RJ, Anderson RM, Johnson SC, Kastman EK, Kosmatka KJ, Beasley TM, et al. Caloric Restriction Delays Disease Onset and Mortality in Rhesus Monkeys. Science 2009;325:201–4. 10.1126/science.1173635.

14 Partridge L, Gems D, Withers DJ. Sex and Death: What Is the Connection? Cell 2005;120:461–72. 10.1016/j.cell.2005.01.026.

15 Piper MDW, Partridge L, Raubenheimer D, Simpson SJ. Dietary restriction and aging: a unifying perspective. Cell Metab 2011;14:154–60. 10.1016/j.cmet.2011.06.013.

16 Simpson SJ, Couteur DGL, Raubenheimer D, Solon-Biet SM, Cooney GJ, Cogger VC, et al. Dietary protein, aging and nutritional geometry. Ageing Res Rev 2017;39:78–86. 10.1016/j.arr.2017.03.001.

17 Holliday R. Food, reproduction and L’ongevity: Is the extended lifespan of calorie-restricted animals an evolutionary adaptation? Bioessays 1989;10:125–7. 10.1002/bies.950100408.

18 Saltiel AR, Kahn CR. Insulin signalling and the regulation of glucose and lipid metabolism. Nature 2001;414:799–806. 10.1038/414799a.

19 Virgilio CD, Loewith R. The TOR signalling network from yeast to man. Int J Biochem Cell Biol 2006;38:1476–81. 10.1016/j.biocel.2006.02.013.

20 Mair W, Piper MDW, Partridge L. Calories Do Not Explain Extension of Life Span by Dietary Restriction in *Drosophila*. Plos Biol 2005;3:e223. 10.1371/journal.pbio.0030223.

21 Grandison RC, Piper MDW, Partridge L. Amino-acid imbalance explains extension of lifespan by dietary restriction in *Drosophila*. Nature 2009;462:1061–4. 10.1038/nature08619.

22 Solon-Biet SM, McMahon AC, Ballard JWO, Ruohonen K, Wu LE, Cogger VC, et al. The Ratio of Macronutrients, Not Caloric Intake, Dictates Cardiometabolic Health, Aging, and Longevity in Ad Libitum-Fed Mice. Cell Metab 2014;19:418–30. 10.1016/j.cmet.2014.02.009.

23 Zanco B, Mirth CK, Sgrò CM, Piper MD. A dietary sterol trade-off determines lifespan responses to dietary restriction in *Drosophila* melanogaster females. Elife 2021;10:e62335. 10.7554/elife.62335.

24 Tavernarakis N. Ageing and the regulation of protein synthesis: a balancing act? Trends Cell Biol 2008;18:228–35. 10.1016/j.tcb.2008.02.004.

25 Babygirija R, Lamming DW. The regulation of healthspan and lifespan by dietary amino acids. Transl Medicine Aging 2021;5:17–30. 10.1016/j.tma.2021.05.001.

26 Green CL, Lamming DW, Fontana L. Molecular mechanisms of dietary restriction promoting health and longevity. Nat Rev Mol Cell Biol 2022;23:56–73. 10.1038/s41580-021-00411-4.

27 Srivastava A, Lu J, Gadalla DS, Hendrich O, Grönke S, Partridge L. The Role of GCN2 Kinase in Mediating the Effects of Amino Acids on Longevity and Feeding Behaviour in *Drosophila*. Frontiers Aging 2022;3:944466. 10.3389/fragi.2022.944466.

28 Sang JH, King RC. Nutritional requirements of axenically cultured *Drosophila* melanogaster adults. J Exp Biol 1961:793–809. 10.1242/jeb.38.4.793.

29 Consuegra J, Grenier T, Baa-Puyoulet P, Rahioui I, Akherraz H, Gervais H, et al. *Drosophila*-associated bacteria differentially shape the nutritional requirements of their host during juvenile growth. Plos Biol 2020;18:e3000681. 10.1371/journal.pbio.3000681.

30 Gallinetti J, Harputlugil E, Mitchell JR. Amino acid sensing in dietary-restriction-mediated longevity: roles of signal-transducing kinases GCN2 and TOR. Biochem J 2012;449:1–10. 10.1042/bj20121098.

31 Dever TE, Feng L, Wek RC, Cigan AM, Donahue TF, Hinnebusch AG. Phosphorylation of initiation factor 2α by protein kinase GCN2 mediates gene-specific translational control of GCN4 in yeast. Cell 1992;68:585–96. 10.1016/0092-8674(92)90193-g.

32 Olsen DS, Jordan B, Chen D, Wek RC, Cavener DR. Isolation of the gene encoding the *Drosophila* melanogaster homolog of the Saccharomyces cerevisiae GCN2 eIF-2alpha kinase. Genetics 1998;149:1495–509. 10.1093/genetics/149.3.1495.

33 Harding HP, Novoa I, Zhang Y, Zeng H, Wek R, Schapira M, et al. Regulated Translation Initiation Controls Stress-Induced Gene Expression in Mammalian Cells. Mol Cell 2000;6:1099–108. 10.1016/s1097-2765(00)00108-8.

34 Harding HP, Zhang Y, Zeng H, Novoa I, Lu PD, Calfon M, et al. An Integrated Stress Response Regulates Amino Acid Metabolism and Resistance to Oxidative Stress. Mol Cell 2003;11:619–33. 10.1016/s1097-2765(03)00105-9.

35 Vattem KM, Wek RC. Reinitiation involving upstream ORFs regulates ATF4 mRNA translation in mammalian cells. P Natl Acad Sci Usa 2004;101:11269–74. 10.1073/pnas.0400541101.

36 Kang M-J, Vasudevan D, Kang K, Kim K, Park J-E, Zhang N, et al. 4E-BP is a target of the GCN2–ATF4 pathway during *Drosophila* development and aging. J Cell Biol 2017;216:115–29. 10.1083/jcb.201511073.

37 Geltosky JE, Mitchell HK. Developmental regulation of phenylalanine hydroxylase activity in *Drosophila* melanogaster. Biochem Genet 1980;18:781–91. 10.1007/bf00484593.

38 Kosakamoto H, Okamoto N, Aikawa H, Sugiura Y, Suematsu M, Niwa R, et al. Sensing of the non-essential amino acid tyrosine governs the response to protein restriction in *Drosophila*. Nat Metabolism 2022;4:944–59. 10.1038/s42255-022-00608-7.

39 Matsuura A, Tsukada M, Wada Y, Ohsumi Y. Apg1p, a novel protein kinase required for the autophagic process in Saccharomyces cerevisiae. Gene 1997;192:245–50. 10.1016/s0378-1119(97)00084-x.

40 Kamada Y, Funakoshi T, Shintani T, Nagano K, Ohsumi M, Ohsumi Y. Tor-Mediated Induction of Autophagy via an Apg1 Protein Kinase Complex. J Cell Biology 2000;150:1507–13. 10.1083/jcb.150.6.1507.

41 Chang Y-Y, Neufeld TP. An Atg1/Atg13 complex with multiple roles in TOR-mediated autophagy regulation. Mol Biol Cell 2009;20:2004–14. 10.1091/mbc.e08-12-1250.

42 Ganley IG, Lam DH, Wang J, Ding X, Chen S, Jiang X. ULK1·ATG13·FIP200 Complex Mediates mTOR Signaling and Is Essential for Autophagy. J Biol Chem 2009;284:12297–305. 10.1074/jbc.m900573200.

43 Hosokawa N, Hara T, Kaizuka T, Kishi C, Takamura A, Miura Y, et al. Nutrient-dependent mTORC1 Association with the ULK1–Atg13–FIP200 Complex Required for Autophagy. Mol Biol Cell 2009;20:1981–91. 10.1091/mbc.e08-12-1248.

44 Jung CH, Jun CB, Ro S-H, Kim Y-M, Otto NM, Cao J, et al. ULK-Atg13-FIP200 Complexes Mediate mTOR Signaling to the Autophagy Machinery. Mol Biol Cell 2009;20:1992–2003. 10.1091/mbc.e08-12-1249.

45 Giannakou ME, Goss M, Jünger MA, Hafen E, Leevers SJ, Partridge L. Long-Lived *Drosophila* with Overexpressed dFOXO in Adult Fat Body. Science 2004;305:361–361. 10.1126/science.1098219.

46 Denton D, Xu T, Dayan S, Nicolson S, Kumar S. Crosstalk between Dpp and Tor signaling coordinates autophagy-dependent midgut degradation. Cell Death Dis 2019;10:111. 10.1038/s41419-019-1368-9.

47 Zinke I, Schütz CS, Katzenberger JD, Bauer M, Pankratz MJ. Nutrient control of gene expression in *Drosophila*: microarray analysis of starvation and sugar-dependent response. Embo J 2002;21:6162–73. 10.1093/emboj/cdf600.

48 Scott RC, Schuldiner O, Neufeld TP. Role and Regulation of Starvation-Induced Autophagy in the *Drosophila* Fat Body. Dev Cell 2004;7:167–78. 10.1016/j.devcel.2004.07.009.

49 Williams GC. Natural Selection, the Costs of Reproduction, and a Refinement of Lack’s Principle. Am Nat 1966;100:687–90. 10.1086/282461.

50 Kirkwood TBL. Evolution of ageing. Nature 1977;270:301–4. 10.1038/270301a0.

51 Shanley DP, Kirkwood TBL. Calorie restriction and aging: a life-history analysis. Evolution 2000;54:740–50. 10.1111/j.0014-3820.2000.tb00076.x.

52 Wek RC, Jackson BM, Hinnebusch AG. Juxtaposition of domains homologous to protein kinases and histidyl-tRNA synthetases in GCN2 protein suggests a mechanism for coupling GCN4 expression to amino acid availability. Proc National Acad Sci 1989;86:4579–83. 10.1073/pnas.86.12.4579.

53 Malzer E, Szajewska-Skuta M, Dalton LE, Thomas SE, Hu N, Skaer H, et al. Coordinate regulation of eIF2α phosphorylation by PPP1R15 and GCN2 is required during *Drosophila* development. J Cell Sci 2013;126:1406–15. 10.1242/jcs.117614.

54 Mizushima N, Komatsu M. Autophagy: Renovation of Cells and Tissues. Cell 2011;147:728–41. 10.1016/j.cell.2011.10.026.

55 Whitney ML, Jefferson LS, Kimball SR. ATF4 is necessary and sufficient for ER stress-induced upregulation of REDD1 expression. Biochem Bioph Res Co 2009;379:451–5. 10.1016/j.bbrc.2008.12.079.

56 Xu D, Dai W, Kutzler L, Lacko HA, Jefferson LS, Dennis MD, et al. ATF4-Mediated Upregulation of REDD1 and Sestrin2 Suppresses mTORC1 Activity during Prolonged Leucine Deprivation. J Nutrition 2019. 10.1093/jn/nxz309.

57 Telfer WH, Kunkel JG. The Function and Evolution of Insect Storage Hexamers. Annu Rev Entomol 1991;36:205–28. 10.1146/annurev.en.36.010191.001225.

58 An JH, Blackwell TK. SKN-1 links C. elegans mesendodermal specification to a conserved oxidative stress response. Genes Dev 2003;17:1882–93. 10.1101/gad.1107803.

59 Tullet JMA, Hertweck M, An JH, Baker J, Hwang JY, Liu S, et al. Direct Inhibition of the Longevity-Promoting Factor SKN-1 by Insulin-like Signaling in C. elegans. Cell 2008;132:1025–38. 10.1016/j.cell.2008.01.030.

60 Linford NJ, Bilgir C, Ro J, Pletcher SD. Measurement of Lifespan in *Drosophila melanogaster*. J Vis Exp 2013. 10.3791/50068.

61 Bass TM, Grandison RC, Wong R, Martinez P, Partridge L, Piper MDW. Optimization of Dietary Restriction Protocols in *Drosophila*. Journals Gerontology Ser 2007;62:1071–81. 10.1093/gerona/62.10.1071.

62 Piper MDW, Blanc E, Leitão-Gonçalves R, Yang M, He X, Linford NJ, et al. A holidic medium for *Drosophila* melanogaster. Nat Methods 2014;11:100–5. 10.1038/nmeth.2731.

63 Piper MDW, Soultoukis GA, Blanc E, Mesaros A, Herbert SL, Juricic P, et al. Matching Dietary Amino Acid Balance to the In Silico-Translated Exome Optimizes Growth and Reproduction without Cost to Lifespan. Cell Metab 2017;25:610–21. 10.1016/j.cmet.2017.02.005.

64 Huang C, Foster SR, Shah AD, Kleifeld O, Canals M, Schittenhelm RB, et al. Phosphoproteomic characterization of the signaling network resulting from activation of the chemokine receptor CCR2. J Biol Chem 2020;295:6518–31. 10.1074/jbc.ra119.012026.

65 Tyanova S, Temu T, Cox J. The MaxQuant computational platform for mass spectrometry-based shotgun proteomics. Nat Protoc 2016;11:2301–19. 10.1038/nprot.2016.136.

66 Cox J, Neuhauser N, Michalski A, Scheltema RA, Olsen JV, Mann M. Andromeda: A Peptide Search Engine Integrated into the MaxQuant Environment. J Proteome Res 2011;10:1794–805. 10.1021/pr101065j.

67 Tyanova S, Temu T, Sinitcyn P, Carlson A, Hein MY, Geiger T, et al. The Perseus computational platform for comprehensive analysis of (prote)omics data. Nat Methods 2016;13:731–40. 10.1038/nmeth.3901.

68 RStudio Team. RStudio: Integrated Development Environment for R. Boston, MA: RStudio, Inc; 2018.

69 Yu G, Wang L-G, Han Y, He Q-Y. clusterProfiler: an R Package for Comparing Biological Themes Among Gene Clusters. Omics J Integr Biology 2012;16:284–7. 10.1089/omi.2011.0118.

70 Gramates LS, Agapite J, Attrill H, Calvi BR, Crosby MA, Santos G dos, et al. FlyBase: a guided tour of highlighted features. Genetics 2022;220:iyac035. 10.1093/genetics/iyac035.

71 Kanehisa M, Goto S. KEGG: Kyoto Encyclopedia of Genes and Genomes. Nucleic Acids Res 2000;28:27–30. 10.1093/nar/28.1.27.

72 Therneau T. A Package for Survival Analysis in R. 2015.

73 Fox J, Weisberg S. An R Companion to Applied Regression. Third. Thousand Oaks CA: Sage; 2019.

74 Lenth RV, Bolker B, Buerkner P, Giné-Vázquez I, Herve M, Jung M, et al. emmeans: Estimated Marginal Means, aka Least-Squares Means. 2023.

