## Supplemental Table 2 for "GCN2 mediates access to stored amino acids for somatic maintenance during *Drosophila* ageing"

**Table S2 – Protein interactions as referenced in Figure 4.**

The details of each interaction were gathered from studies using yeast (*Saccharomyces cerevisiae*), flies (*Drosophila melanogaster),* murine *(Mus musculus*) cell culture or human (*Homo sapiens)* cell culture.

| **Interaction (*Drosophila* gene orthologues in brackets)** | **Characterised in** | **References** |
|---|---|---|
| Insufficient AAs activate GCN2 (*GCN2)* | *S. cerevisiae*<br>*D. melanogaster*<br>*H. sapiens* cell culture | [1] Wek, Jackson and Hinnebusch, 1989<br>[2] Olsen *et al.*, 1998<br>[3] Harding *et al.*, 2000 |
| GCN2 (*GCN2)* activates eIF2α (*eIF2α)* | *S. cerevisiae*<br>*D. melanogaster* | [4] Dever *et al.*, 1992<br>[5] Malzer *et al.*, 2013 |
| eIF2α (*eIF2α)* activates ATF4 (*crc)* | *D. melanogaster* cell culture<br>*M. musculus* cell culture | [6] Kang *et al.*, 2017<br>[7] Vattem and Wek, 2004 |
| ATF4 (*crc)* activates REDD1 (*scyl*, *chrb*) | *M. musculus* cell culture<br>*D. melanogaster* | [8] Kosakamoto *et al.* 2022<br>[9] Whitney, Jefferson and Kimball, 2009<br>[10] Xu *et al.*, 2019 |
| REDD1 (*scyl*, *chrb*) inhibits mTORC1 (*Tor*, *raptor* + others) | *D. melanogaster*<br>*M. musculus* cell culture<br>*H. sapiens* cell culture | [11] Kosakamoto *et al.* 2022<br>[12] Reiling and Hafen, 2004<br>[13] Brugarolas *et al.*, 2003<br>[14] Corradetti, Inoki and Guan, 2005<br>[15] DeYoung *et al.*, 2008<br>[16] Garami *et al.*, 2003<br>[17] Tee *et al.*, 2003<br>[18] Li *et al.*, 2004)<br>[19] Sofer *et al.*, 2005 |
| mTORC1 (*Tor*, *raptor* + others) inhibits ATG1/ATG13 (*Atg1, Atg13)* complex formation | *S. cerevisiae*<br>*D. melanogaster*<br>*M. musculus* cell culture<br>*H. sapiens* cell culture | [20] Kamada *et al.*, 2000<br>[21] Matsuura *et al.*, 1997<br>[22] Chang and Neufeld, 2009<br>[23] Ganley *et al.*, 2009<br>[24] Hosokawa *et al.*, 2009<br>[25] Jung *et al.*, 2009 |
